# Hsp70 chaperone blocks α-synuclein oligomer formation via a novel engagement mechanism

**DOI:** 10.1101/2020.05.15.098699

**Authors:** Jiahui Tao, Amandine Berthet, Rose Citron, Robert Stanley, Jason Gestwicki, David A. Agard, Lisa McConlogue

**Affiliations:** Department of Biochemistry and Biophysics, University of California San Francisco, San Francisco, CA 94143, USA; Gladstone Institute of Neurological Disease, San Francisco, CA 94158, USA; Department of Pharmaceutical Chemistry, Institute for Neurodegenerative Diseases and UCSF Weill Institute for Neurosciences, University of California San Francisco, San Francisco, CA 94143, USA

**Keywords:** α-Synuclein, chaperone, Hsp70, protein misfolding, Parkinson’s Disease, Lewy body dementia, synucleinopathy

## Abstract

Over-expression and aggregation of α-synuclein (ASyn) are linked to the onset and pathology of Parkinson’s disease and related synucleinopathies. Elevated levels of the stress induced chaperone, Hsp70, protects against ASyn misfolding and ASyn-driven neurodegeneration in cell and animal models, yet there is minimal mechanistic understanding of this important protective pathway. It is generally assumed that Hsp70 binds to ASyn using its canonical and promiscuous substrate-binding cleft to limit aggregation. Here we report that this activity is due to a novel and unexpected mode of Hsp70 action, involving neither ATP nor the typical substrate-binding cleft. We use novel ASyn oligomerization assays to show that Hsp70 directly blocks ASyn oligomerization, an early event in ASyn misfolding. Using truncations, mutations and inhibitors, we confirmed that Hsp70 interacts with ASyn via an as yet unidentified, non-canonical interaction site in the C-terminal domain. Finally, a biological role for a non-canonical interaction was observed in H4 neuroglioma cells. Together, these findings suggest that new chemical approaches will be required to target Hsp70-ASyn interaction in synucleinopathies. Such approaches are likely to be more specific than targeting Hsp70 canonical actions. Additionally, these results raise the question of whether other misfolded proteins might also engage via the same non-canonical mechanism.

## Introduction

Neuropathological, biochemical and genetic evidence strongly implicates α-Synuclein (ASyn) in the onset and progression of Parkinson’s disease (PD) and related syncucleinopathies including Lewy body dementias (LBD), multiple systems atrophy and Alzheimer’s disease. Aggregated ASyn inclusions are a hallmark of these diseases[1], and ASyn gene mutations or multiplications cause early onset PD or LBD[2, 3, 4]. The sequential misfolding of ASyn into oligomers and fibrils is central to the pathogenesis of the synucleinopathies. ASyn gene multiplications and mutations causing PD and LBD are associated with enhanced oligomer formation[5, 6, 7]. Significant evidence supports the hypothesis that prion-like transmission of misfolded ASyn underlies disease progression[8, 9]. The severity of disease correlates with the progressive spread of aggregated ASyn in patients[10], and misfolding is associated with toxicity in cell and animal models[11]. As shown for other amyloid misfolding proteins, compelling evidence supports pre-fibrillar ASyn oligomers and not fibrillar deposits as the pathogenic species in disease[5, 12, 13, 14, 15, 16]. ASyn oligomers are directly toxic to cells[16], and mutations enhancing ASyn oligomer formation increase ASyn toxicity in neurons and rodents[12, 13, 14].

The constitutive (Hsc70) and inducible (Hsp70) forms of the 70-KDa cytosolic heat shock molecular chaperones (Hsp70s) assist a wide variety of folding processes and provide broad protection against protein misfolding in the cell. Hsp70 polymorphisms are associated with PD[17] and in PD patients stress induced Hsp70 accumulates in a thwarted attempt to clear aggregated ASyn[18]. Although Hsp70 is protective against ASyn pathogenicity in cell and animal models[19, 20, 21, 22, 23, 24], little is known about its protective mechanisms. The general biochemical mechanisms of Hsp70 action are well understood. Hsp70s are composed of a nucleotide-binding domain (NBD), which has ATPase activity, and a substrate binding domain (SBD), which contains the canonical binding cleft for misfolded proteins. In addition, Hsp70s have disordered, C-terminal regions that engage in additional protein-protein interactions (PPIs). Hsp70 has two known mechanisms of mitigating protein misfolding: an ATP-cycling dependent “foldase” action which is able to restore damaged proteins and an ATP-independent “holdase” mode which binds to unfolded proteins to prevent aggregation. Biochemical investigations demonstrate that Hsp70s block ASyn fibrillization[25, 26, 27, 28, 29] in an ATP-independent manner[26, 27, 30] indicating a “holdase” based mechanism. It was assumed that this holdase-like activity was mediated via Hsp70’s canonical substrate binding site in the SBD, which is required for other known holdase roles[31]. The promiscuous engagement of a broad array of Hsp70 substrates with the canonical substrate binding site has precluded effective attempts at therapeutic targeting of Hsp70 canonical actions for neurodegenerative diseases.

Due to the importance of ASyn in disease, we directly investigated the mechanism of Hsp70’s activities in the early stages of Asyn oligomerization. Using novel ASyn oligomerization biochemical assays, we confirmed that Hsp70 directly blocks ASyn oligomerization in an ATP-independent holdase manner. However, we unexpectedly found that Hsp70’s activity was not mediated by the canonical Hsp70 substrate binding site. Recently, a few examples of non-canonical Hsp70 interactions have been discovered[32], but we further found that these non-canonical sites are not involved. Hsp70 blocking of ASyn oligomer formation in cells is mediated by a similar non-canonical mechanism. Together, these results indicating the presence of a previously unidentified non-canonical interaction site. These findings have major implications for the use of Hsp70 as a drug target in synucleinopathies.

## Results

### Novel ASyn oligomerization assays

Because fibrillization assays are complex and oligomers are the likely pathologic species, we reasoned that biochemical investigation of Hsp70’s action on the earliest stages of ASyn oligomer formation would be most informative. In prior studies, we established biochemical ASyn oligomerization assays that are based on either the complementation of split Gaussia luciferase (gLuc) or FRET between fluorophores on separate ASyn molecules[33]. Here we develop an improved biochemical complementation assay using split NanoBit luciferase (nLuc)[34] tags placed on separate ASyn molecules (Fig 1). Importantly, unlike either split fluorescent proteins (such as split GFP) or the Gaussia split luciferase[35], the nLuc complementation system employs fully reversible and quite weakly interacting components, and its components are stable and non-linkable via disulfide bonds[34]. Purified ASyn fused at its C-terminus with either the large (LgBit) or the small (smBit) portion of nLuc were incubated, and the formation of oligomers monitored by reconstituted nLuc activity. Split nLuc tags placed on separate ASyn molecules reconstituted luciferase activity, whereas removal of ASyn from one of the tags gave minimal background signal (Fig 1A). As expected for an early stage misfolding event, the formation of oligomers did not require the agitation needed for the formation of thioflavin T (ThT) reactive fibrillar species nor have the characteristic lag phase seen in ASyn fibrillization reactions (Fig 1B). The oligomerization reaction also showed a strict dependence on temperature, with efficient formation observed at 37°C and negligible formation at room or lower temperatures (Fig 1C). The highly quantitative split nLuc assay also allowed us to monitor the ASyn concentration dependence of the reaction (Fig 1D), providing insight into the number of ASyn monomers that must assemble to reconstitute luciferase activity (Fig 1E). The data showed that the minimal assembly required to detect a nLuc signal was between dimers or trimers (n= 2.6; Fig 1E), which matched well with that seen in the absence of complementation tags using a FRET based assay (n=2.4; Fig 1F). Note however, that nLuc signal developed at lower ASyn concentrations than the FRET signal, indicating the inherent avidity expected with any complementation system (Fig 1E vs 1F). The increased sensitivity is expected to be beneficial when exploring the effects of oligomerization modulators.

**Figure 1.**
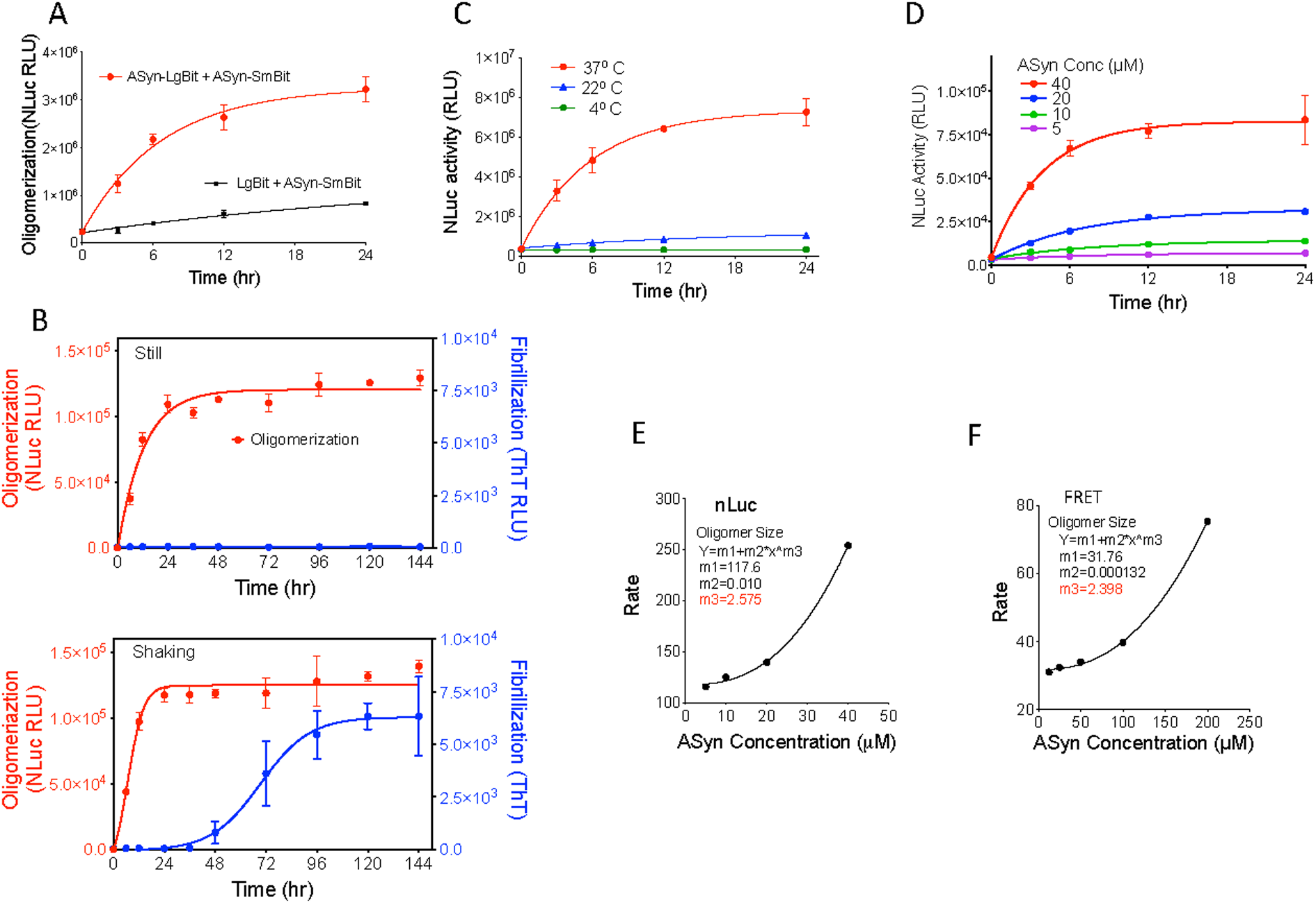
ASyn biochemical oligomerization assays. **A)** Split nLuc tagged ASyn biochemical oligomerization assay development. LgBiT protein alone (black) or ASyn protein tagged with the LgBit (ASyn-LgBiT, red) are mixed with ASyn protein tagged with the SmBit (ASyn-SmBiT) at 10 μM each, incubated without shaking at 37°C, and assayed for nLuc activity at various times. Oligomerization is detected by complementation of the split tags attached to ASyn. **B)** 10 μM each of ASyn-LgBiT and ASyn-SmBiT are incubated in PBS under still (top panel) or shaking (bottom panel) conditions and analyzed for either ASyn oligomerization using the nLuc assay (red) or fibrilization via ThT fluorescence (blue). Oligomerization did not require shaking whereas fibrilization did. **C)** Temperature dependence of split nLuc tagged ASyn oligomerization assay. Tagged ASyn was incubated at various temperatures and nLuc activity measured. 37°C was required for robust oligomer formation. **D)** Various concentrations of ASyn-LgBiT and ASyn-SmBiT were assayed for ASyn oligomerization using the nLuc assay. Oligomerization is dependent on ASyn concentration. **E)** Initial rates of ASyn oligomerization in the nLuc assay, calculated from the data in D, versus ASyn concentration are shown and analyzed by the formula Y=m1+m2*x^m3 to derive a minimal size of detected ASyn oligomer (m3) of 2.6 ASyn molecules. **F)** Initial rates of ASyn oligomerization in the FRET assay, calculated from the data in figure S4, versus ASyn concentration are shown. Analysis as in E derived a minimal size of detected ASyn oligomer (m3) of 2.4 ASyn molecules. In all graphs the symbols show mean and bars show ± s.d., n=3.

Combined size exclusion chromatography and multi angle light scattering (SEC-MALS) analyses of either nLuc tagged (Fig 2A) or untagged ASyn (Fig 2B) species indicated that ASyn converts from a largely monomeric to a largely trimeric state over the incubation time course of the assays (24 hours for nLuc tagged species, 48 hours for untagged FRET assay species[33]), indicating that we are monitoring the early stages of oligomerization. Previously, it was shown that oligomers capable of seeding fibril formation can be formed in *in vitro* biochemical assays[36]. To test that our complementation tags do not interfere with fibril seeding ability, we took samples at different time points from our nLuc oligomerization assay and added it to an excess of untagged monomeric ASyn. This mixture was incubated with shaking at 37°C and fibril formation monitored with ThT fluorescence. The oligomers formed in reaction conditions without shaking, which precludes fibril formation (Fig 1B), were able to seed fibril formation in a time (Fig 2C) and concentration-dependent manner (Figs 2D, E) and thus contained oligomers on pathway for fibril formation.

**Figure 2.**
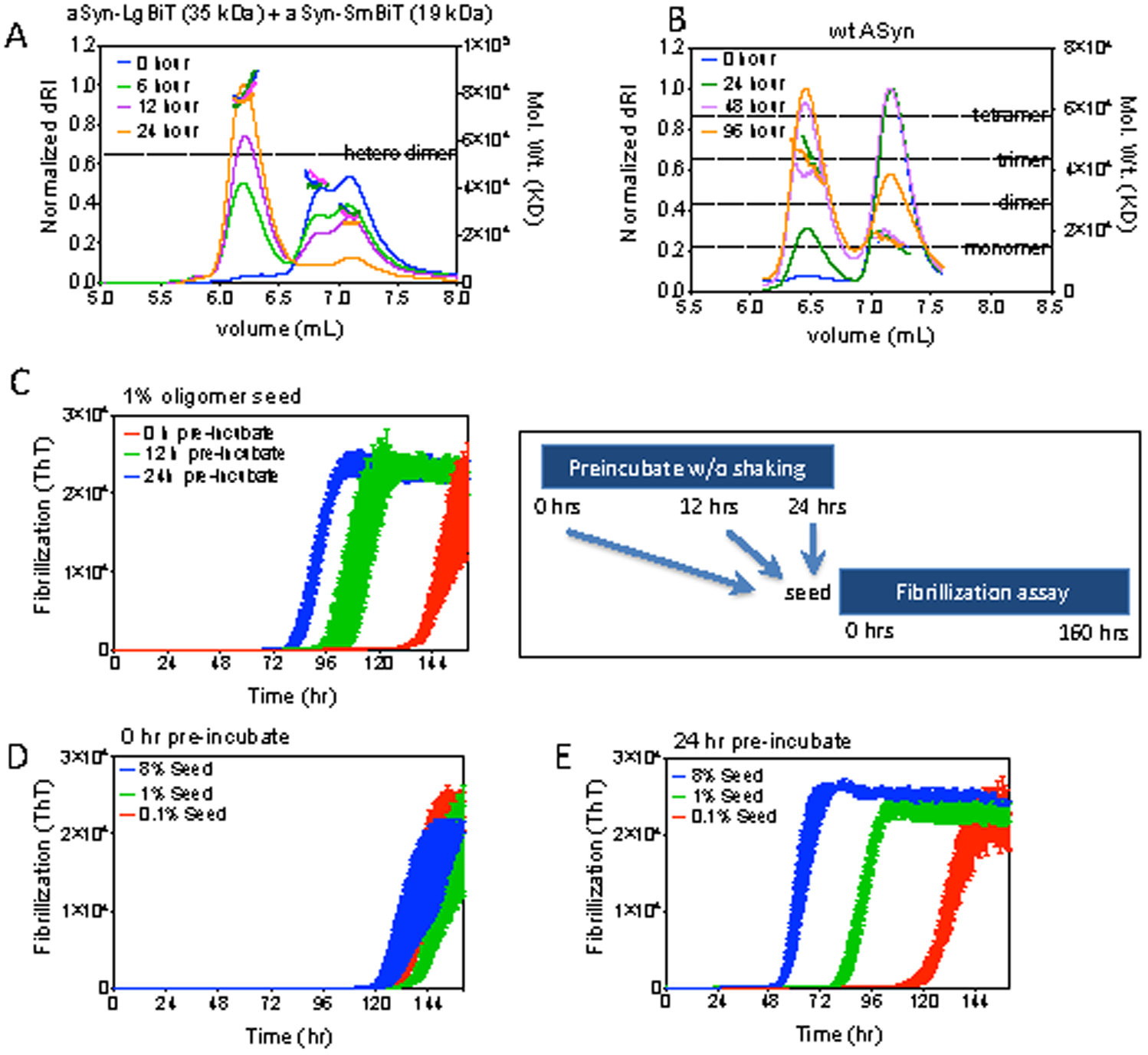
ASyn-Nluc Oligomers can seed ASyn fibrillization. 50 µM each of ASyn-LgBiT and ASyn-SmBiT **(A)** or 100 µM untagged ASyn **(B)** were incubated for the indicated times at 37°C and then analyzed by size exclusion chromatography with multi-angle light scattering (SEC-MALS). Solid lines are refractive index and dotted lines are the calculated molecular weights. ASyn converts from a largely monomeric to a largely trimeric state over the incubation time course of our assays (24 hr. nLuc, 96 hr. FRET). **C) D) E)** Pre-formed split nLuc tagged ASyn oligomers are used as seeds for an ASyn fibrillization assay as described in methods. Nluc-tagged ASyn oligomers are formed in a still preincubation of ASyn-LgBiT and ASyn-SmBiT and subsequently seeded into an untagged ASyn fibrillization assay. Various percentages of ASyn-Nluc oligomers in a large excess of untagged ASyn are incubated as described in methods and ASyn fibrils formed are quantified using ThT fluorencence. **C)** seeding capabilities increase with preincubation time for a 1% seeding. **D)** 0 time of preincubated does not seed fibrillization. **E)** Seeding capacity of 24 hour preincubated split nLuc tagged ASyn increases with amount of seed added. Mean ± SEM shown, n=3.

### Hsp70 blocks ASyn oligomerization in an ATP independent manner

We next tested Hsp70 for impact on ASyn oligomerization. In three ASyn oligomerization assays, split Gaussia luciferase (gLuc)[33](Fig 3A), nLuc (Fig 3B), and FRET[33] (Fig 3C), Hsp70 impairs oligomer formation at similar IC_50_ concentrations (11.6 μM gLuc, 9.1 μM nLuc, 19.3 μM FRET). Hsp70 is unable to disassemble preformed oligomers (Fig 3D), which is consistent with previous findings[29].

**Figure 3.**
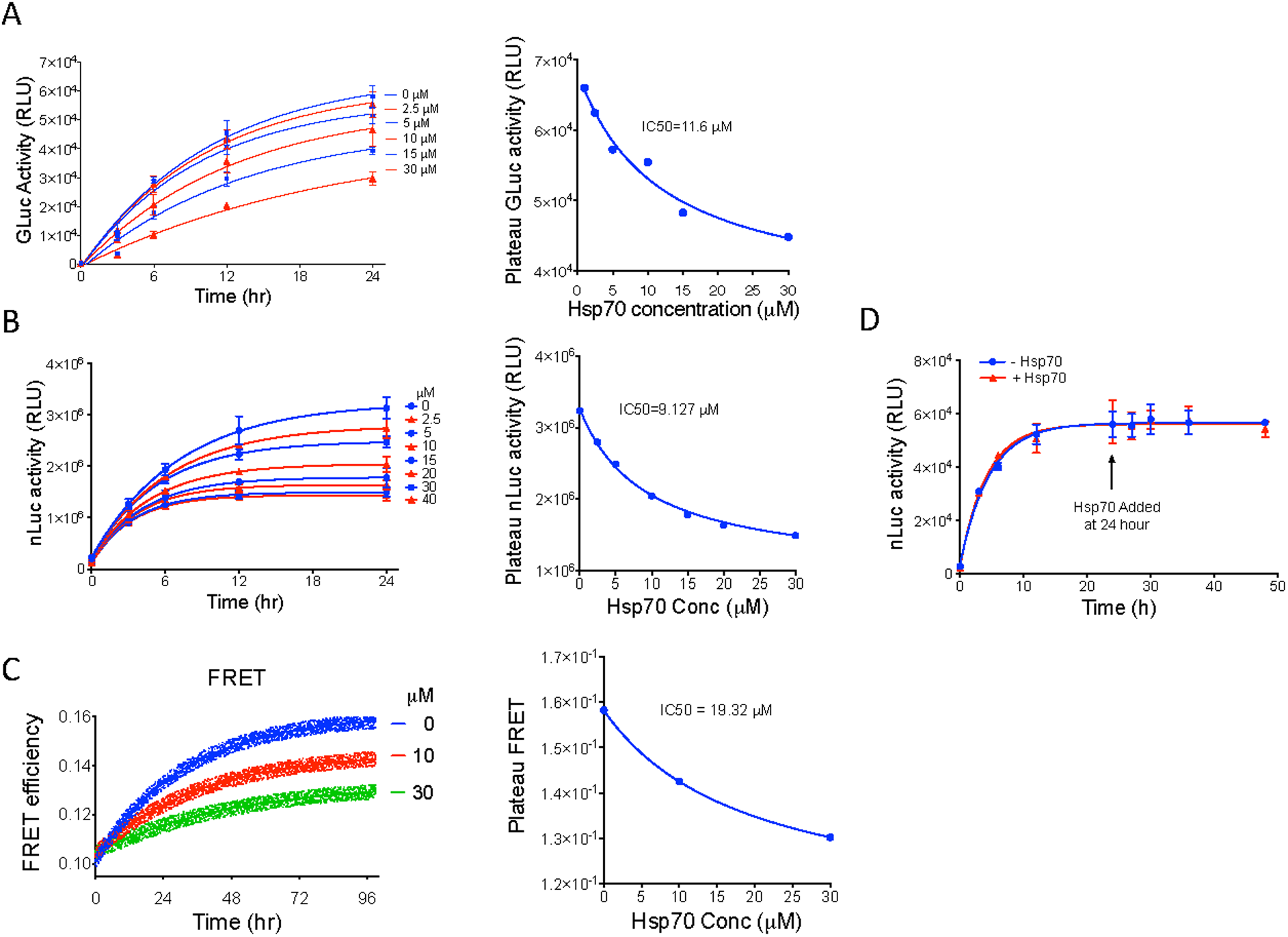
Hsp70 impairs ASyn oligomerization in multiple assays. Various concentrations of Hsp70 were added to the **A)** split gLuc[33], **B)** split nLuc and **C)** FRET[33] based ASyn oligomerization assays. All assays were as in methods. **B)** 5 μM each of ASyn-SmBiT and ASyn-LgBiT were incubated with 1 mM ADP, 1 mM Mg. **C)** ASyn-Cy3 and ASyn-Cy5 were incubated with 0 (blue) 10 (red) or 30 (green) μM Hsp70 with 1 mM MgCl2, 1 mM ADP. Triplicate replicates (circles) are shown with first order kinetic fit (line). Hsp70 impaired ASyn oligomerization in all three assays. The IC_50_ of Hsp70 for each assay is calculated based on the plateau signal at 24 hours using the GraphPad Prism nonlinear regression model for dose response (three parameters). **D)** 10 μM of each of ASyn-SmBiT and ASyn-LgBiT were incubated for 24 hours after which 20 μM Hsp70, 1 mM Mg and 1 mM ADP were added, and the reactions further incubated. Hsp70 fails to disassemble already formed ASyn oligomers. Means ± s.d. shown, n=3.

It is well appreciated that Hsp70s engage substrates via a substrate binding pocket in the SBD. In turn, the affinity of substrates for this cleft is regulated by the nucleotide state of the NBD, with weak affinity in the ATP-bound state and tight binding in the ADP-bound state[31] (Fig 4A). In the ATP state, the N-terminal NBD (green in Fig 4A) interacts with both parts of the C-terminal SBD (yellow and orange in Fig 4A), sequestering the α-helical lid subdomain (orange in Fig 4A) away from the bottom side of the substrate-binding cleft (yellow in Fig 4A). In Figure 4B, the location of the canonical substrate binding pocket, present in the ADP bound form, is highlighted by the presence of a bound peptide fragment. Thus, in the canonical model, ATP binding and hydrolysis drive cycles of substrate binding and release[37, 38].

**Figure 4.**
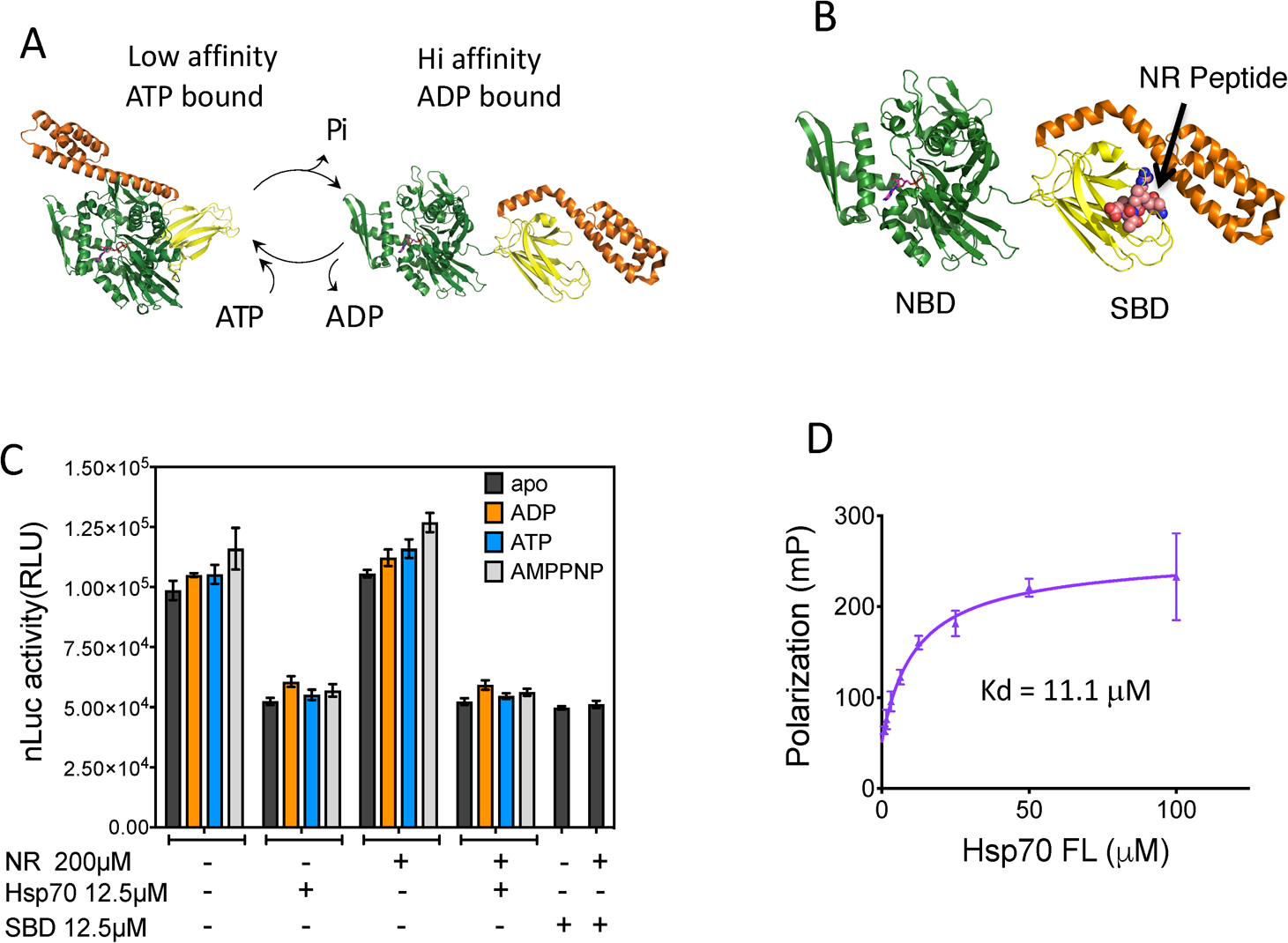
Hsp70 blockage of ASyn oligomerization requires neither ATP cycling nor binding to the canonical substrate binding site. **A)** Schematic of canonical action of Hsp70 family members. Structures are based on the E. Coli Hsp70 homologue, DnaK. ATP hydrolysis drives Hsp70 from a low (ATP) to a high affinity (ADP) substrate binding state. Nucleotides bind to the N terminal nucleotide binding domain (NBD, green, residues 1-388). Substrate binds to a pocket formed in the C-terminal substrate binding domain (SBD, Yellow/orange residues 389-602) by the lid region (orange, residues 508-602) folding over the β-sheet core of the SBD (yellow, residues 389-507) in the ADP bound conformation. Protein data bank identifiers are 2kho, 4b9q and 1dkx. **B)** Structure of NR peptide bound to the E.Coli Hsp70 homologue DnaK is shown. NR peptide binds to the canonical substrate binding site. **C)** 10 μM each of ASyn-LgBiT and ASyn-SmBiT, with or without full length Hsp70, Hsp70-SBD (see Fig 5), 2 mM nucleotide and/or the NR peptide as indicated were incubated in 384-well plates for 24 hours and nLuc luciferase activity measured as described in methods. ADP stabilizes the substrate bound conformation, and AMPNP, a non-hydrolyzable ATP analogue, stabilizes the ATP bound state. Neither nucleotide, NR nor truncation impacted Hsp70 blockage of ASyn oligomerization. Means ± s.d. shown, n=3. **D)** Binding of NR peptide to Hsp70 was measured by fluorescence polarization of N-terminal 5-FAM (Fluorescein) label on the NR peptide as described in methods. NR binds with a Kd of 11.1 μM. Means ± s.d. shown, n=3.

By contrast with this normal mode of action, previous studies have shown that both Hsp70 and Hsc70 can impair ASyn fibrillization in the absence of ATP cycling and even without the NBD[26, 27, 30]. We wanted to both assess whether Hsp70 is working in a similar manner to block oligomerization and to better understand this unexpected mechanism. We thus analyzed the nucleotide dependence of Hsp70’s ability to block ASyn oligomerization in our assay (Fig 4C). We found that neither the addition of ATP nor conditions that blocked ATP cycling (addition of ADP or the non-hydrolyzable ATP analogue AMPNP) impacted Hsp70’s ability to block oligomerization (Fig 4C). The NBD was also not required, as the SBD alone had activity equivalent to the full length Hsp70 (Figs 4C and 5A). Taken together, these data are consistent with a holdase model of Hsp70 action, in which the Hsp70 SBD is able to sequester ASyn monomers, blocking their ability to oligomerize or form fibrils.

### Hsp70 does not engage ASyn via its canonical substrate binding site

It has generally been assumed that Hsp70 engages ASyn via its canonical substrate binding site. To determine if this were the case, we tested the ability of a well-characterized Hsp70 substrate, the NR peptide, to compete with ASyn binding, thereby restoring the ability of ASyn to oligomerize. As determined using a fluorescent polarization assay, NR peptide binds Hsp70 with a Kd of 11.1 μM (Fig 4D) yet does not inhibit Hsp70’s action on ASyn oligomerization at concentrations up to 200 μM (Fig 4C). This result was unexpected, and it indicates that the part of the SBD required to block ASyn oligomerization is distinct from the canonical substrate binding pocket. Further, the NR peptide had no impact on the activity of full-length Hsp70 regardless of nucleotide state (Fig 4C), indicating that the non-canonical ASyn binding site must be fully available independent of Hsp70 domain organization or occupancy of the canonical substrate binding pocket.

To define what part(s) of the SBD was most relevant for inhibiting ASyn oligomerization, we examined the activity of additional Hsp70 truncations. We focused these studies on removing parts of Hsp70 that are known to interact with co-chaperones or engage in other protein-protein interactions (PPIs). Specifically, co-chaperones are known to bind the linker between the NBD-SBD and to EEVD residues at the extreme C-terminus of the lid domain[32]. We found that a version of Hsp70’s NBD, which includes the inter-domain linker, was still inactive (Fig 5A), suggesting that this PPI site was not involved. The EEVD sequence mediates Hsp70 engagement both with Hsp90 which promotes refolding, and with CHIP which engages proteasomal degradation[32]. Deleting the C-terminal EEVD residues did not alter the ability to block ASyn oligomerization *in vitro* (delEEVD Fig 5B), showing that this known PPI site is not responsible. However, we found that deleting the entire lid partially impaired the activity of the SBD (Fig 5A), suggesting that this subdomain, outside the EEVD, could play a role. Finally, we introduced a point mutant, K589A, which is sufficient to block binding of Hsp70 to endosomes[32, 39], but found that it is also not relevant for ASyn *in vitro* (K589A, Fig 5B). Together these results suggest that a unique and unmapped surface(s) on the lid domain, and perhaps the core β-sheet domain, are required for productive engagement of Hsp70 with ASyn.

**Figure 5.**
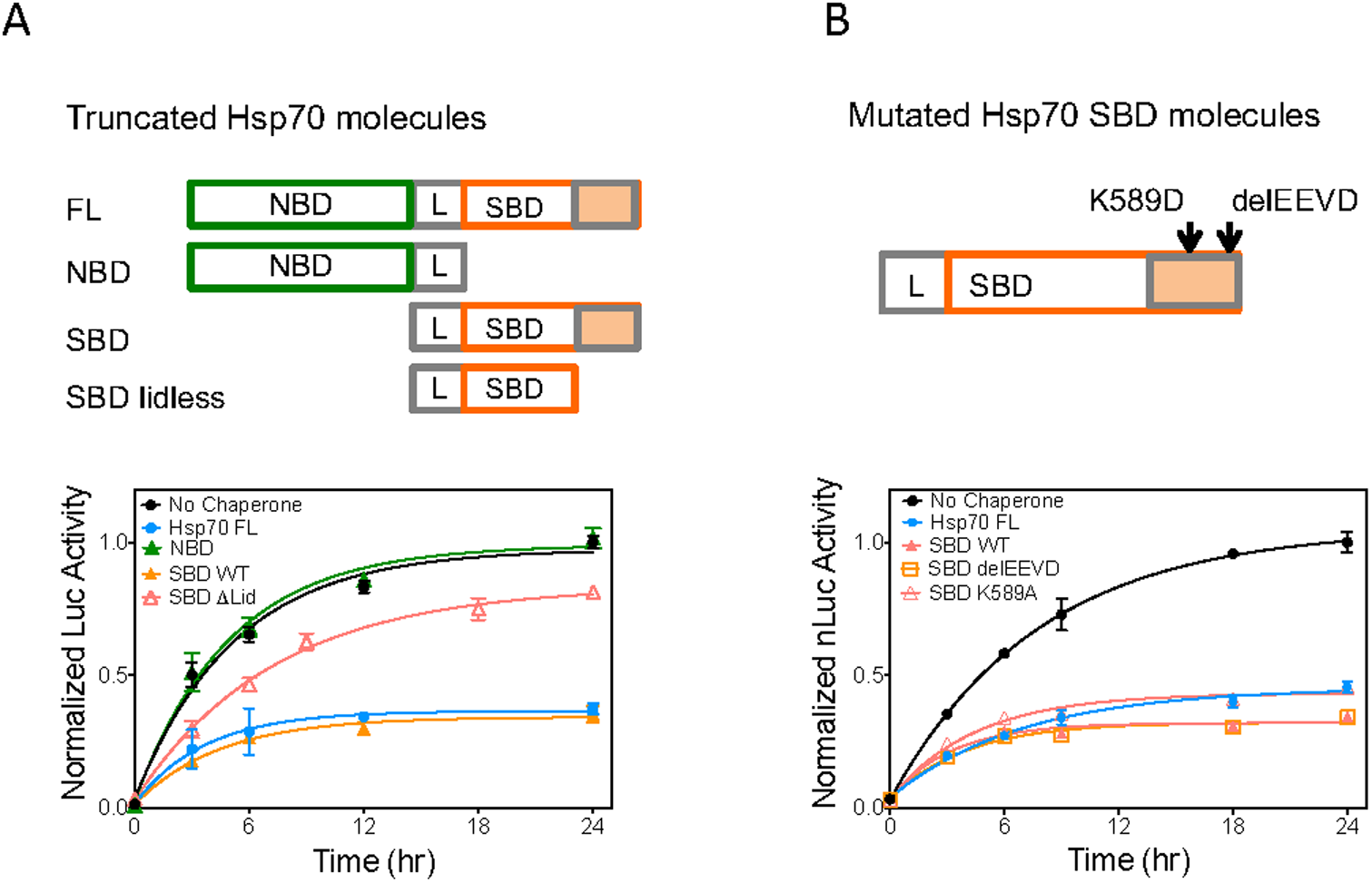
Activity of truncated and mutated Hsp70 species. **A)** Activity of truncated Hsp70 species. **Top:** Schematic of Hsp70 truncations. Nucleotide binding domain (NBD, green, residues 1-381), linker region (L, grey, residues 382-393), substrate binding domain (SBD, orange open and filled, residues 394-641), SBD without the lid region (SBD Δlid, orange open, residues 394-510). **Bottom:** Hsp70 fragments are tested for ability to impair ASyn oligomerization. 10 μM each of ASyn-LgBiT and ASyn-SmBiT were incubated with 10 μM of the indicated Hsp70 and Nluc activity measured as described in methods. Means ± s.d. shown, n=3. **B)** Activity of mutated Hsp70 species. **Top:** Schematic of Hsp70 mutations in the Hsp70 SBD. K589A blocks Hsp70 recognition of phosphotidylserine modified proteins and subsequent engagement for endosomal autophagy. C-terminal EEVD sequence deleted blocks Hsp70 engagement both with Hsp90 which promotes refolding, and with CHIP which engages proteasomal degradation. **Bottom:** Hsp70 fragments are tested for ability to impair ASyn oligomerization. ASyn oligomerization assays were performed as in A. Hsp70 impairment of ASyn oligomerization does not depend on Hsp70 interaction sites used endosomal autophagy nor for Hsp90/CHIP engagement. Means ± s.d. shown, n=3.

### A similar mechanism of Hsp70 action blocks ASyn oligomerization in cells

We next extended these biochemical observations to H4 neuroglioma cells to understand if the non-canonical interaction was relevant in cells. We found that the previously reported cell model using overexpression of split gLuc tags on ASyn[21] drives ASyn into large S129 phospho-synuclein positive aggregates in H4 cells (Supplemental Information and Fig S1). We considered that the presence of these large aggregates could complicate analyses of oligomerization using the split gLuc tagged system, so we established an alternative, quantitative cellular ASyn oligomerization assay using the split nLuc complementation system which does not form such aggregates (Fig S1). We established conditions for both cell handling and assay parameters in which nLuc activity resulting from split nLuc tag complementation was proportional to the amount of tagged ASyn transiently overexpressed for both cellular and media compartments (Supplemental Information and Fig S2). This assay provided an alternative cell-based platform for studying the interaction of Hsp70 with ASyn.

Using this assay, we found that over-expression of either full-length Hsp70 or the Hsp70 SBD reduced ASyn oligomers in H4 neuroglioma cells (Fig 6A). Reproducibility of expression of transfected ASyn-LgBit and of ASyn-SmBit in each of the samples was confirmed by Western analyses (Fig 6B). Neither the wildtype nor mutant SBD constructs used here showed any indication of toxicity or perturbation of proteostasis, as evidenced by the unchanged cell numbers (Fig S3) and the consistent levels of actin and markers of the stress responses (Hsp27and endogenous Hsp70) (Fig 6B). Unexpectedly, the transfected Hsp70 runs slightly higher than the endogenous human H4 cell Hsp70. The smaller endogenous band is unlikely to result from cross reaction with Hsc70 or BiP, as these Hsp70 homologues are both larger than Hsp70. Rather, this effect could be due to expression of a different human Hsp70 variant in the H4 neuroglioma cell line under basal conditions. For example, the sequence we chose to overexpress was based on the consensus entries for Hsp70A1 (GenBank NM_005345.6 and BC002453.2). However, there is an alternative GenBank entry (M11717.1), which differs by two amino acid changes and an additional one deleted. More likely it is the Hsp70 variant HspA2 which is two amino acids smaller than the Hsp70A1 version we overexpressed[40].

**Figure 6.**
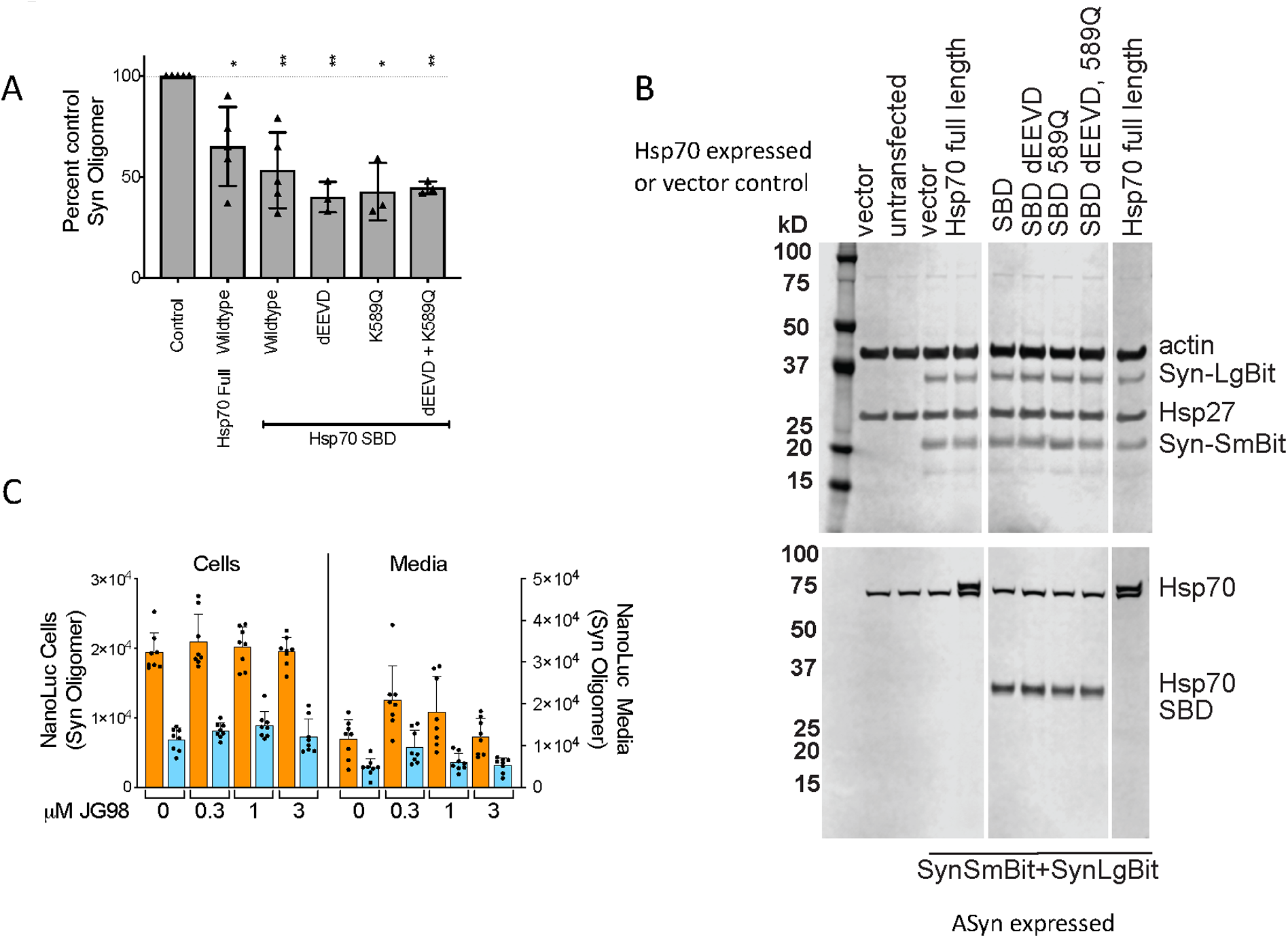
Neither ATPase activity nor downstream engagements are required for Hsp70-mediated inhibition of oligomerization in H4 neuroglioma cells. **A)** The impact of overexpression of full length and truncated Hsp70 without or with mutations blocking downstream pathways is tested on ASyn oligomer formation in H4 cells transiently transfected with vectors co-expressing ASyn-LgBiT and ASyn-SmBiT. Samples normalized to split NanoBit tagged ASyn (ASyn-SmBit+ASyn-LgBit) control without Hsp70 are compiled from multiple experiments. Each symbol is the mean of 6 to 12 wells of a separate experiment and error bars are ± s.d. Statistics are one sample t-test two-tailed for difference from control (100), * p< 0.05, **<0.01. All Hsp70 variants tested impaired ASyn oligomer formation in H4 cells. **B)** Western analyses of expressed and cellular proteins from samples run in parallel to panel A. Expression of transfected proteins is robust and consistent and does not impact levels of actin, endogenous Hsp70 nor the stress protein Hsp27 indicating that overexpression of mutant Hsp70 does not perturb cellular proteostasis. **C)** vector (orange) or full length Hsp70 (cyan) are co-transfected with vectors expressing ASyn-LgBiT or ASyn-SmBiT and treated with 0.1% DMSO alone or containing allosteric Hsp70 ATPase inhibitor JG98 at non-toxic doses that are ≥3-fold higher than the previously reported EC_50_s[44]. JG-98 was toxic at 10 μM. Fugene transfection was as described in Supplemental Information. ASyn oligomers were measured by assaying H4 cells and media at 48 and 24 hours respectively post transfection for reconstituted nLuc activity. Hsp70 ATPase inhibition by JG98 had no impact on its ability to block ASyn oligomerization in cells. Mean ± s.d. with individual wells shown, n=8.

Next, we wanted to further test the role of a non-canonical interaction. First, treatment with JG-98, an allosteric Hsp70 ATPase inhibitor that impairs Hsp70’s canonical action in other pathways[41, 42, 43, 44], did not block Hsp70’s ability to reduce ASyn oligomers in H4 cells or media (Fig 6C). Additionally, mutations in the SBD, such as the EEVD deletion and the endosome blocking K589Q mutation[32, 39], did not block the ability of the SBD to reduce ASyn oligomer levels in cells (Figs 6A). As was the case for the SBD, none of these mutations showed toxicity or perturbation of proteostasis (Fig 6B and S3). Thus, like we observed in the biochemical system, Hsp70 SBD does not act to block ASyn oligomerization through canonical or previously identified, non-canonical SBD binding sites. Rather, this important interaction appears to occur through a new non-canonical site, which involves parts of the SBD and lid.

## Discussion

Although Hsp70 is protective against ASyn pathogenicity in cell and animal models[19, 20, 21, 22, 23, 24], little is known about its protective mechanisms. Biochemical investigations demonstrate that the Hsp70s block ASyn fibrillization[25, 26, 27, 28, 29] largely via an ATP-independent holdase mechanism[26, 27, 30]. Due to the multi-step complexity and variability of fibrillization assays, the stage at which this inhibition occurs is not established. In particular, Hsp70’s impact on ASyn oligomerization, the initial step of fibrillization, has not been directly determined[25, 26, 27, 28, 30]. Using both previously reported assays[33] and a novel, biochemical ASyn oligomerization approach, we show that Hsp70 directly blocks ASyn oligomerization in a nucleotide-independent manner. Surprisingly, we show that a competitive inhibitor of the Hsp70 canonical substrate binding site, the NR peptide, does not block this action, indicating that a non-canonical site must be responsible for retarding ASyn oligomerization. Thus, it may be possible to separate Hsp70’s blocking of ASyn oligomerization from its action on other necessary cellular substrates.

Recently, a few examples of Hsp70 interactions distinct from the canonical substrate binding site have been discovered driving Hsp70 engagement with XIAP substrates[45] and downstream autophagic and co-chaperone actions[32]. As the XIAP substrates bind to the NBD of Hsp70[45] and ASyn oligomerization blocking activity lies within the SBD, they cannot share an Hsp70 engagement site. Deletion of the C-terminal EEVD motif of Hsp70 which engages Hsp90 to coordinate protein refolding as well as CHIP to mediate transfer of proteins to the proteasome[32], has no impact on Hsp70 blocking ASyn oligomerization. Hsp70 mutation K589Q which blocks Hsp70 directed transfer of phosphytidyl serine modified proteins to endosomal autophagy[32, 39], also has no impact on ASyn oligomerization. Thus Hsp70’s ability to suppress ASyn oligomerization relies on a previously unidentified non-canonical interaction site. Interestingly, the Hsp70 region responsible for chaperone mediated autophagy is as yet unidentified, so could not be directly tested and it remains a possibility.

In order to probe the mechanism of Hsp70 suppression of ASyn oligomerization in H4 neuroglioma cells, we established a new cellular ASyn oligomerization assay based on complementation of split nLuc tags placed on ASyn. In our hands, split gLuc tags drive ASyn into large aggregates possibly confounding analyses of oligomerization. The split nLuc tagged system has the advantage of very low intrinsic binding of the split tags and furthermore did not drive ASyn aggregation. Using this novel assay, we show that Hsp70 suppression of ASyn is mediated by an ATP-independent mechanism, as it is driven entirely by the SBD and is insensitive to Hsp70 ATPase inhibition. Furthermore, as was the case in our biochemical experiments, it does not rely upon previously mapped non canonical Hsp70 engagement regions. Thus, a similar mechanism of Hsp70 suppression of ASyn oligomerization is at play in cells.

Because ASyn oligomers are likely the most pathogenic misfolded ASyn species[5, 46], we hypothesize that the novel non-canonical Hsp70 site responsible for blocking ASyn oligomerization also provides protection against ASyn pathogenicity in animal models and in disease (Fig 7). Further biochemical and structural mechanistic studies will be needed to determine the exact molecular mechanisms of Hsp70 suppression of ASyn oligomerization.

**Figure 7.**
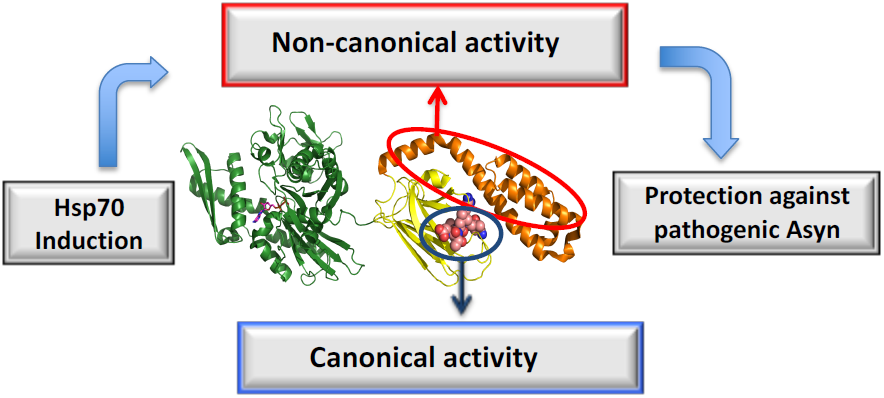
Hypothesis: Novel engagement of Hsp70 with ASyn via a site separate from the canonical substrate binding site leads to specific and protective action on ASyn. We propose a novel mechanism in which a non-canonical alternative binding site for ASyn on Hsp70 is responsible for both blocking ASyn oligomerization and for conferring protection against ASyn pathogenicity.

Such structural studies will allow targeted mutagenesis of Hsp70 ASyn engagement sites to be tested in *in vivo* studies for protection against neurodegeneration to directly test our hypothesis.

The discovery that Hsp70 blocks ASyn oligomerization in a novel manner makes it possible to develop therapeutic approaches that enhance these beneficial effects without interfering with Hsp70’s many cellular functions. An underlying assumption of the field has been that protective Hsp70 actions on ASyn all occur via its canonical substrate binding site. An important caveat of this assumption is that a chemical inhibitor of Hsp70’s ATP cycling or canonical interactions would influence all of Hsp70’s many roles in the cell. This scenario might be expected to result in untoward effects complicating therapeutic application for neurodegenerative diseases. One possible therapeutic approach targeting Hsp70’s non-canonical action on ASyn would be the development of targeted ASyn pharmacological chaperones[33, 47] which are small molecules that bind to and stabilize select ASyn conformations. Such compounds could impact ASyn’s mode of interaction with Hsp70 allowing for enhanced blocking of ASyn oligomer formation. In support of this proposed approach, we have reported the development of small molecule pharmacological chaperones that engage ASyn and is so doing reverse multiple different ASyn malfunctions[33, 48]. Alternatively, it is possible that allosteric modulating sites exist on Hsp70 whose engagement by small molecules could enhance this desirable activity. Hsp70 is a complex molecule with significant allosteric modulation by co-factors. Small molecules activators of canonical Hsp70 activity have been identified[49]. Hsp70 has significant structural complexity with rich opportunity for regulatory pockets, including in its SBD lid region supporting the possibility of finding such activating molecules.

In conclusion, these studies indicate that Hsp70 engages ASyn to prevent oligomer formation in a novel manner distinct from its normal canonical substrate binding site. This implies that specifically targeting Hsp70 to ASyn for treatment of PD, LBD and related synucleinopathies is possible. A high priority will be to determine the exact molecular mechanism of this novel activity.

## Methods

Additional Methods are available in Supplemental Information

### Molecular cloning

#### Bacterial expression vectors

Gaussia luciferase (pET28b-αSyn-GLuc1 and pET28b-αSyn-GLuc2) and FRET (pET28b-aSyn-Q99C) vectors were generated as described[33]. NanoBit luciferase (nLuc) was split into two fragments: LgBiT (N-terminal 158 residues); SmBiT (C-terminal 11 residues)[34]. Either LgBiT or SmBiT was fused to the C terminus of human ASyn with a flexible linker (GGGGSGGGGSSG) placed between ASyn and the nLuc tags. The bacterial expression vectors, pET28b-αSyn-LgBiT and pET28b-αSyn-SmBiT were constructed by inserting either coding regions into pET28b vector (Novagen, Darmstadt, Germany) via NcoI/NotI sites in the multiple cloning site. The human Hsp70 coding region (HSPA11A, Genecopia, amino acid sequence identical to GenBank: BC002453.2) was amplified using Q5 High-Fidelity DNA Polymerase (New England Biolabs, Ipswich MA, USA) and inserted into the TOPO cloning site in pET151/D-TOPO E. Coli expression vector (Invitrogen, Waltham MA, USA). The pET151/D-TOPO-Hsp70 plasmid was used for recombinant expression of Hsp70 protein. Truncated Hsp70 constructs were made using a Q5 site-directed mutagenesis kit (New England Biolabs, Ipswich MA, USA) according to the manufacturer’s instructions. All plasmids were sequence verified.

#### Mammalian expression vectors

pMKHsp70-1A: The human Hsp70 coding region (Hsp70 gene 1A, GenBank: BC002453.2, and NM_005345) from the 2072 b.p. NheI-XbaI fragment of pcDNA3.1-Hsp70 (gift from Dr. Chad Dickey) was combined with the 7035 bp XbaI-NheI fragment of expression vector pMK1252 (gift from Dr. Martin Kampman, UCSF). pVLHsp70-1A: The human Hsp70 coding region (Hsp70 gene 1A, GenBank: BC002453.2, and NM_005345.6) from the 1991 bp BamHI-EcoRI(filled in) fragment of pcDNA3.1-Hsp70 was cloned into the BamHI/HpaI polylinker sites of pLV-Bobi[50]. The inserted fragment and flanking vector regions for all constructs were sequenced verified using at least 4 independent runs covering both strands. pLV-Bobi[50], an empty expression vector and pLV-αSyn vector (LV-αSyn)[51] expressing untagged ASyn were gifts from Dr. Brian Spencer of UCSD.

### E. Coli protein expression and purification

BL21(DE3) *E. coli* cells were transformed with desired plasmids and cultured in LB media at 37°C. At OD_600_ 0.6-0.8, the culture temperature was lowered to 20°C, IPTG (GoldBio, St. Louis, MO, USA) was added at 0.5 mM to induce protein expression and cultures further incubated at 20°C overnight. The cells were harvested by centrifugation at 4,000 g for 20 minutes in an Avanti J-26 XPI centrifuge with a JLA 8.1000 rotor (Beckman Coulter, Brea, CA, USA).

#### Proteins αSyn-GLuc1, αSyn-GLuc2, αSyn-LgBiT, Hsp70

Plasmid transformed E. coli cells were resuspended in 25 mM Tris, 500 mM NaCl, 0.5 mM TCEP, pH 8.0 and then lysed in the presence of EDTA-free Protease Inhibitor Cocktail (Roche Life Sciences, Penzberg, Germany) using an EmulsiFlex-C3 (Avestin, Ottawa, Canada). The lysate was cleared by centrifugation at 30,000 g for 30 minutes in an JA 25.50 rotor (Beckman Coulter, Brea, CA, USA). The His-tagged target protein was purified by Ni-NTA gravity-flow chromatography (Qiagen, Hilden, Germany). The eluted protein was subjected to MonoQ 10/100 GL (GE Healthcare Lifesciences, Marlborough, MA, USA) chromatography using a 0-600 mM NaCl gradient elution. αSyn-GLuc1 and αSyn-GLuc2 were polished by HiLoad 16/60 Superdex 75 size exclusion chromatography (SEC) (GE Healthcare Lifesciences, Marlborough, MA, USA), collecting fractions eluting at 47-61ml. αSyn-LgBiT and Hsp70 were polished by HiLoad 16/60 Superdex 200 SEC (GE Healthcare Lifesciences, Marlborough, MA, USA); fractions eluting at 68-82 ml and 78-86 ml were collected respectively. The harvested protein was concentrated by Amicon Ultra-15 centrifugal filter units (Millipore Sigma, Burlington, MA, USA), filtered through a 0.22 µm filter (E&K Scientific, Santa Clara, CA, USA), flash frozen and stored in aliquots at -80 °C.

#### Proteins αSyn-SmBiT, αSyn-Q99C

Plasmid transformed E. coli cells were resuspended in 20 mM Tris, pH 8.0 and lysed by boiling for 30 minutes in the presence of an EDTA-free Protease Inhibitor Cocktail (Roche Life Sciences, Penzberg, Germany). The lysate was cleared by centrifugation at 30,000 g for 30 minutes in a JA 25.50 rotor (Beckman Coulter, Brea, CA, USA). Streptomycin sulfate was added to the lysate at 10 mg/ml to precipitate DNA. After a 30 minute incubation at 4°C, the lysate was cleared by centrifugation at 30,000 g for 30 minutes. Ammonium sulfate was added to the lysate to at 0.36 g/ml to precipitate protein. After incubation at 4°C overnight, the protein was pelleted by centrifugation at 30,000 g for 30 minutes. The protein was resuspended in 20 mM Tris, 1 mM DTT, pH 8.0, then subjected to MonoQ 10/100 GL (GE Healthcare Lifesciences, Marlborough, MA, USA) chromatography using a 0-600 mM NaCl gradient elution. αSyn-SmBiT and αSyn-Q99C were polished by HiLoad 16/60 Superdex 200 SEC (GE Healthcare Lifesciences, Marlborough, MA, USA), collecting fractions eluted at 78-86 ml. The harvested protein was concentrated, filtered and stored as described above.

### Biochemical ASyn oligomerization assays

The split Gaussia luciferase (gLuc) based oligomerization assay was performed as described[33], and used for Fig 3A, with the exception that 1 mM ADP, 1 mM MgCl2 and Hsp70 at various concentrations were added at the beginning of the assay. The Forster resonance energy transfer (FRET) oligomerization assay was carried out as described[33], except that 1 mM ADP, 1 mM MgCl2 and Hsp70 at various concentrations were added in Fig 3C. For the split NanoLuc luciferase (nLuc) oligomerization assay, different concentrations of αSyn-LgBiT and αSyn-SmBiT were mixed at a 1:1 molar ratio in PBS (pH 7.4). The mixtures were incubated at 37°C in micro PCR tubes on a thermal cycler (BioRad DNA Engine PTC-200). At the indicated time points, 10 µl of the mixture were placed in a single well of a black/clear bottom 384-well microplate (Greiner, Kremsmünster, Austria). Oligomerization was quantified by measuring the luminescence on a SpectraMax L Microplate Reader (Molecular Devices, Aliso Viejo, CA, USA) using either an h-CTZ protocol or a NanoFuel protocol. For the h-CTZ protocol, 40 µl of h-CTZ (40 µM, NanoLight Technology, Norman, OK, USA) was injected into the wells with the samples using the injector integrated in the SpectraMax L. The luminescence was measured with 2 second delay and 3 second integration times. The h-CTZ protocol applies to figures 1A, 1C, 1D, 3B, 3D, 4C, 5A and 5B. Alternatively, for the Nanofuel protocol, NanoFuel GLOW Assay kit (NanoLight Technology, Norman, OK, USA) reagent was added into each well according to manufacturer’s instructions. The luminescence measurements were performed on the SpectraMax L with 8 minute delay and 1 second integration times. This protocol was used for figures 1B and 4C. In experiments used to evaluate Hsp70 impact, 1 mM nucleotide (ATP, ADP or AMPNP) and 1 mM MgCl2 and Hsp70 or Hsp70 mutants at different concentrations were added as shown at the indicated time points (Figs 3B, 3D, 4C 5A, 5B). For all three assays 1 μM Ydj1, an hsp40 which stimulates ATP cycling and presents clients to Hsp70, was added in some cases (Figs 3A, 3B, 3C, 3D) until it was found to have no impact.

### ASyn fibrillization Assays

To measure fibrillization, ASyn proteins were incubated with or without shaking at 1000 rpm on a ThermoMixer (Eppendorf 5350, Hauppauge, NY, USA) at 37°C as indicated. The dye Thioflavin T (ThT) is widely used to detect amyloid fibrillization as it fluoresces upon binding to fibrils[52]. Samples were diluted 20-fold into 25 µM Thioflavin in PBS and incubated at room temperature for 15 minutes to 1 hour in 384 plates. ThT fluorescence was read on a SpectraMax M5 microplate reader (excitation 440 nm, emission 482 nm) in endpoint mode. This protocol applies to Fig 1B. For experiments measuring seeding of ASyn fibrillization (Fig 2 C, D and E), ASyn-Nluc oligomers were formed by incubating 10 μM each of ASyn-LgBiT and ASyn-SmBiT for 0, 12, or 24 hours at 37°C in a thermal cycler (BioRad DNA Engine PTC-200, Hercules, CA, USA). The pre-formed oligomers were used as seeds for an ASyn fibrillization assay. Untagged monomeric ASyn was seeded with 8%, 1% or 0.1% (molar ratio) of the seeds to a final ASyn protein concentration of 200 μM and incubated at 37°C with shaking in 384-well microplate with clear bottom (Greiner, Kremsmünster, Austria) sealed with AlumaSeal II (Hampton Research, Aliso Viejo, CA, USA) to prevent evaporation. ASyn fibril formation was quantified by reading ThT fluorescence on a SpectraMax M5 microplate reader (excitation 440 nm, emission 482 nm) in kinetic mode over the time course of fibrillization.

### Size-exclusion chromatography coupled to multi-angled light scattering analysis (SEC-MALS)

For αSyn-LgBiT/αSyn-SmBiT oligomers, recombinantly expressed and purified αSyn-LgBiT and αSyn-SmBiT were mixed at a 1:1 molar ratio in PBS to a final concentration of 50 µM each. The mixture was incubated at 37°C in PCR tubes on a thermal cycler (BioRad DNA Engine PTC-200). After various times 50 µl of the mixture was taken and subject to SEC using a Shodex KW-802.5 column (HICHROM, Reading, UK) pre-equilibrated in PBS. The SEC was coupled to a static 18-angle light scattering detector (DAWN HELEOS-II, Wyatt Technology, Goleta, CA, USA), a UV detector, and a refractive index detector (Optilab T-rEX, Wyatt Technology, Goleta, CA). Data collection was done at a flow rate of 0.35 ml/min. Data was processed using the analysis package as part of the program ASTRA, yielding the reported molar masses of the protein complexes. For wt ASyn oligomers, recombinantly expressed and purified wt ASyn was diluted to 100 µM in PBS. The protein was incubated at 37°C in PCR tubes on a thermal cycler. At 0, 24, 48 or 96 hours post incubation, 50 µl of samples were taken and analyzed by SEC-MALS. The data collection and processing were carried out following the same protocol for αSyn-LgBiT/αSyn-SmBiT.

### Binding of NR peptide to Hsp70

NR peptide was synthesized with a N-terminal 5-FAM modification by GenScript (Piscataway, NJ, USA). Hsp70 was titrated in triplicate while the NR-peptide concentration remained constant at 20nM. Experiments were performed in filtered PBS with 1mM MgCl and 1mM ADP. Fluorescence polarization of the NR peptide was measured on a SpectraMax M5 plate reader (Molecular Devices, San Jose, CA, USA) with excitation and emission wavelengths of 485nm and 538nm respectively at 37°C. The binding curve was plotted using Prism 6.2 and fit with a single site binding equation curve to obtain a Kd.

### Western Analyses

Western analyses were run as described[33] with the following changes: Purified mouse anti-ASyn antibody clone 42 (BD Biosciences, San Jose, CA) was diluted at 1/1000, rabbit anti Hsp70 antibody (Enzo Life Science, Inc., Farmingdale, NY, ADI-SPA-812) and mouse monoclonal antibody G3.1 anti Hsp27 antibody (Enzo Life Science, Inc., Farmingdale, NY) were diluted at 1/1000, and mouse anti actin antibody clone AC15, (Sigma-Aldrich, St. Louis, MO) was diluted at 1/35,000. Membranes were then washed 4 times for 10 minutes each in PBS 0.1% Tween (PBST) and incubated with secondary antibody donkey anti-mouse infrared 680 and goat anti-rabbit infrared 800 (LI-COR, Lincoln, NE) diluted at 1/10 000 in Odyssey PBS Blocking Buffer (LI-COR, Inc. Lincoln, NE, USA) with 0.2% Tween 20 followed by washing 4 times for 10 minutes each in PBST. Membranes were scanned and quantitated using the Odyssey CLx Imaging System (LI-COR, Lincoln, NE, USA). Hsp70 was visualized in the 800 nm fluorescent channel and actin, αSyn, Hsp27 and the molecular weight markers visualized at 700nm. As previously described[33] proteins are cross-linked to blots by 30 minute treatment with 0.4% PFA in PBS prior to staining to enhance ASyn binding[53] to ensure visualization of all ASyn.

### Cellular nLuc assays

H4 neuroglioma cells (HTB-148; ATCC, Manassas, VA, USA) were passaged in DME containing 10% fetal calf serum (FCS). The cells were plated into polyD-Lysine coated clear bottom white well 96 well plates at 7×10^3^ cells per well for the oligomerization assays and into 6 well plate at 1.5×10^5^ for Western blotting. The following day cells were transfected using either JetPrime (Polyplus, Illkirch-Graffenstaden, France) or Fugene (Promega, Madison, WI, USA) according to the manufacturer’s directions. For Fig 6A and 6B DNA and JetPrime were used at a ratio of reagent to DNA of 2:1. A ratio of plasmids ASyn-Smbit to ASyn-Lgbit at 6:1 of were mixed with either Hsp70-expressing vectors or with control vector not expressing protein (pLV-Bobi) with ratio of ASyn plasmids to Hsp70 expressing or control plasmid of 3:7. After 4 hours, the DNA containing media was removed and cells were fed fresh DME media containing 10% FCS. The following day, the media was changed to Opti-MEM without phenol red (Opti-MEM) with penicillin and streptomycin media. The next day 80 µl of cellular media was transferred into a 96 well white plate for luciferase measurements. To control for possible impacts of protein overexpression on cell numbers, the cells were incubated with Opti-MEM containing Hoechst 33342 (Invitrogen, Carlsbad, CA, USA) diluted at 1/5000 for 30 minutes. The plates were read before and after adding Hoescht on a microplate reader SpectraMax M5 (Molecular Devices, San Jose, CA, USA) with an excitation and emission wavelengths of 350nm and 490nm respectively. The media was removed and replaced with 100 µl of Opti-MEM and cells assayed for luciferase activity. Fugene (Fig 6C) transfection was carried out with a 4:1 ratio of reagent to DNA. 24 hours after transfection the cellular media was replaced with Opti-MEM containing 0.1% DMSO alone or with drugs and incubated a further 24 hours. 80 µl of cellular media was removed for luciferase activity measures, the cells were washed in PBS and 100 µl of Opti-MEM placed in the well and cells assayed for luciferase activity.

Luciferase activity was quantified by measuring the luminescence on a SpectraMax L Microplate Reader (Molecular Devices, San Jose, CA, USA). Briefly, 80 µl for media or 100 µl for cells of h-CTZ (40 µM, NanoLight Technology, Norman, OK, USA) in Opti-Mem was injected into the wells using the injector integrated in the SpectraMax L. The luminescence was measured with 15 second delay and 5 second integration times. The impact of compounds on H4 cell toxicity was assayed using the CytoTox-Glo kit (Promega, Madison, WI).

The synthesis and characterization of compound JG-98 was previously described[44]. The impact of compounds on H4 cell toxicity was assayed using the CytoTox-Glo kit (Promega, Madison, WI).

### Statistical Analysis Methods

Statistical analyses were run using GraphPad Prism software as described. Multiple samples were compared using one-way ANOVA with Dunnett’s and an alpha of 0.05. In figure 6 one sample t-test two-tailed for difference from control (100) was used. General practice significance nomenclature was used (0.1234 (ns), 0.0332 (*), 0.0021 (**), 0.0002(***), <0.0001(****) unless otherwise indicated.

All mandatory laboratory health and safety procedures have been complied with in the course of conducting the experimental work reported herein.

## Acknowledgements

We wish to thank Dr. Brian Spencer for plasmids pLV-Bobi and PLVSyn, Dr. Chad Dickey for plasmid pcDNA3.1-Hsp70, Dr. Martin Kampman for plasmid pMK1252 and Dr. Pamela McLean for plasmids Syn-Luc1 (S1) and Syn-Luc2 (S2) (also referred to as syn-hGLuc(1) and syn-hGLuc(2)[24] respectively. We also thank Pamela McLean for advice on developing the cellular oligomerization assay. We further thank the Gladstone Assay Development and Drug Discovery Core and Dr. Anke Meyer-Franke for advice and use of the ArrayScan automated imaging system. We are particularly grateful to the late T. Gary Rogers for his generous encouragement and to the Rogers Family Foundation for their support of this work (LM, DAA). Additional support was provided by the Howard Hughes Medical Institute (DAA) and the National Institute of Neurological Disorders and Stroke of the National Institutes of Health under award number R21NS092897(LM).

## Author Contributions

Designed and performed research, analyzed data on biochemical oligomerization experiments (J.T., R.C., D.A.A. L.M.). Conceptual contributions on biochemical oligomerization experiments (J.G.). Designed and performed research, analyzed data on cellular oligomerization experiments (A.B., R.S., L.M.), wrote the paper (L.M., J.G., D.A.A.) designed research, supervised and analyzed data on all aspects (L.M., D.A.A.)

## Data and Materials Availability Statement

All plasmid constructs are available with Material Transfer agreement from UCSF and or the J. David Gladstone Institutes.

