## Supplemental Information for "Hsp70 chaperone blocks α-synuclein oligomer formation via a novel engagement mechanism"


### Results

Accumulation of ASyn phosphorylated at position 129 is the major ASyn modification found in Lewy bodies from Lewy Body Disease patients[1] and accumulation of this modification is used to track ASyn aggregate formation in cell and animal models. We found that the previously reported ASyn oligomerization cell model using overexpression of split gLuc tags on ASyn[2] drives ASyn into large S129 phospho-ASyn positive aggregates in H4 neuroglioma cells whereas untagged ASyn or ASyn tagged with Turquoise fluorescent protein does not (Fig S1). As the presence of such aggregates could complicate investigation of oligomers in cells, we tested the split NanoBit[3](nLuc) tags on ASyn designed for our biochemical assay described in the main text for the formation of $PO_4$-ASyn aggregates. ASyn fused at its C-terminus with either the large (LgBit) or the small (smBit) portion of nLuc or a mixture of the two formed few, if any ASyn $PO_4$ positive aggregates (Fig S1). This enabled us to establish a quantitative cellular ASyn oligomerization assay using the split nLuc tagged ASyn system. Split nLuc tags placed on separate ASyn molecules reconstituted luciferase activity, whereas removal of ASyn from one of the tags gave minimal background signal (Fig S2). We established conditions in which nLuc activity resulting from complementation of split nLuc tags attached to ASyn is proportional to the amount of tagged ASyn transiently overexpressed in H4 neuroglioma cells in both cellular and media compartments (Fig S2). Establishing a proportional assay required optimal nLuc substrate type and concentration, optimal transfection conditions, use of optimum medium and optimal delay and integration times in reading the luciferase activities. In order to use Hsp70 variants to probe mechanism we need to show they do not artificially perturb cells. We show that overexpression of the Hsp70 variants and the split nLuc tagged ASyn did not impact cell densities (Fig S3).

### Figures and Legends

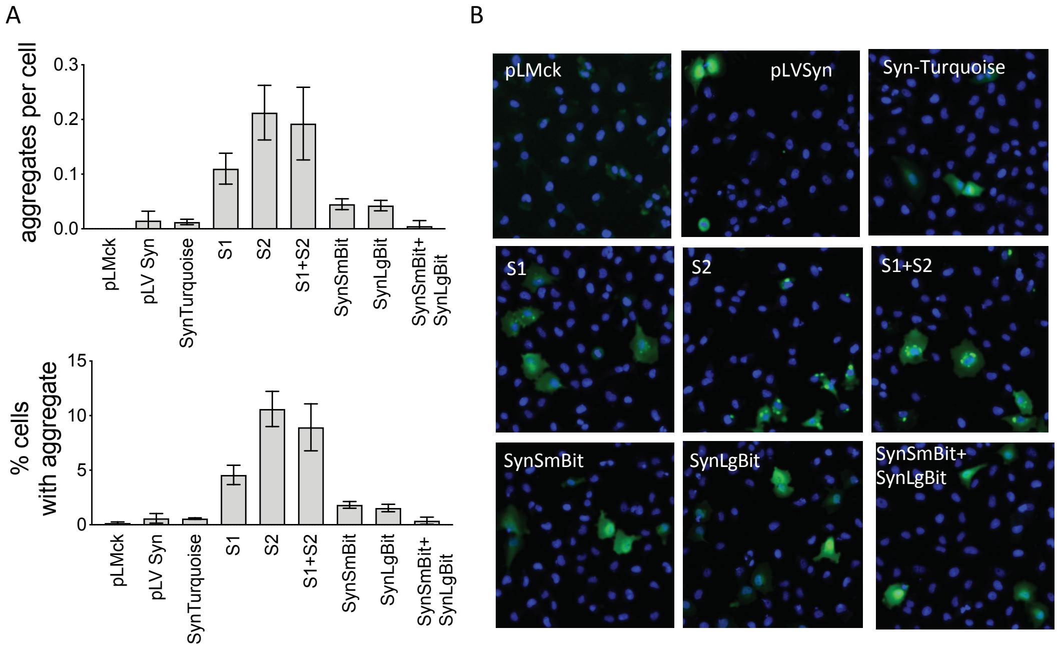


**Figure S1. Split Gaussia but not split nLuc tags on ASyn drive S129 $PO_4$ positive aggregates**. H4 cells were transfected with the indicated plasmids, fixed and stained, and aggregates containing S129 phosphorylated ASyn ($PO_4$-ASyn) quantitated on an Arrayscan automated imaging system as described in methods. A) Quantitation of aggregates per cell (top) and the percentage of cells with aggregates (bottom) indicates that expressing split Gaussia luciferase tagged ASyn (S1 and S2) generates abundant aggregates, whereas vectors expressing nLuc split tags (SynSmBit and SynLgBit) have few to none. Control empty vector (pLMck), untagged ASyn (pLVSyn) and ASyn tagged with the fluorescent protein Turquoise (plasmid pCKSynT1, SynTurquoise) have no aggregates. Means ±s.d., n=4 wells. B) Representative images from each plasmid with nuclei (blue) and $PO_4$-ASyn (green) shown.

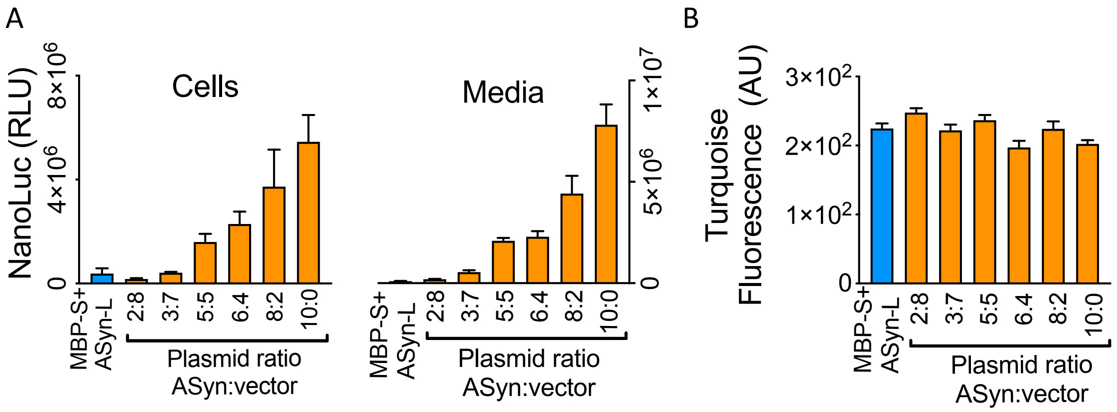


**Figure S2. Development of a cellular ASyn oligomerization assay.**
Control H4 cells were transfected with equal amounts of maltose binding protein fused with nLuc small bit (MBP-S, plasmid pckMBP-LgBit) and ASyn fused to nLuc large Bit (ASyn-L, plasmid pckSyn-LgBit) (blue). An equal mixture of plasmids pckSyn-SmBit (ASyn-S) and pckSyn-LgBit (ASyn-L) were diluted with non-expressing vector (orange) as indicated and transiently transfected into H4 cells as described in methods. The 10:0 ratio is equivalent to the amounts of split nLuc tags in the control (blue) samples. A trace (1/10) amount of control vector expressing turquoise fluorescent protein (pCKSynT1) was included for each sample to measure overall transfection efficiency. 48 hours post transfection cells were assayed for turquoise fluorescence and cells and media were assayed for nLuc activity as described in methods. nLuc activity is dependent on presence of a mixture of both tagged forms of ASyn. The amount of nLuc measured is proportional to the amount of ASyn plasmid transfected for both cells and media. Turquoise controls indicate reproducible transfection efficiencies between samples. Means ± s.d. n= 3.

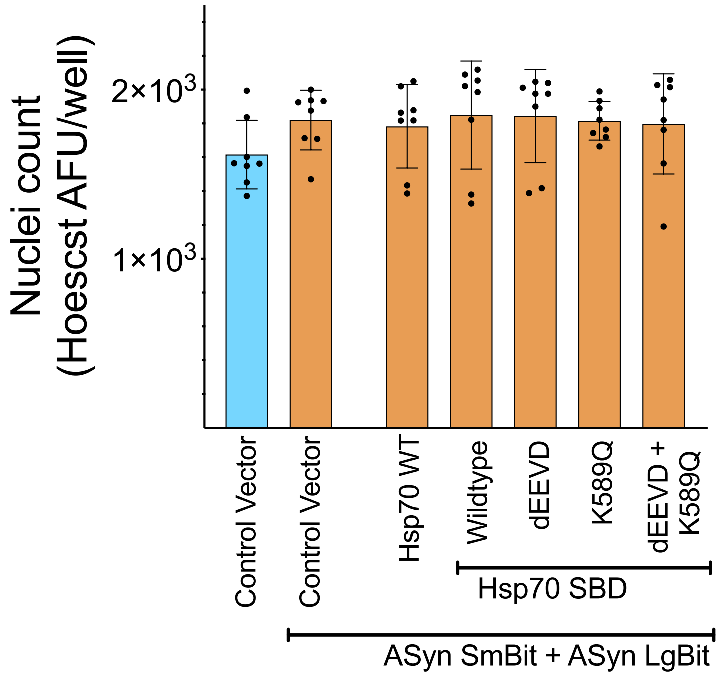


**S3. Lack of toxicity of transfected ASyn and Hsp70 constructs.**
H4 cells were transfected with control non-expressing pVLBobi vector (blue), with a mixture of plasmids pckSyn-SmBit (ASyn-SmBit) and pckSyn-LgBit (ASyn-LgBit) (orange) and with the indicated variant of Hsp70 (Hsp70 WT is full length wildtype) as described for JetPrime transfection in the methods of the main paper. Two days later the nuclei are stained with Hoechst dye and Hoechst fluorescence quantitated as a surrogate for cell number. Overexpression of none of the Hsp70 variants showed impact on cell densities. Means ± s.d. n= 3.

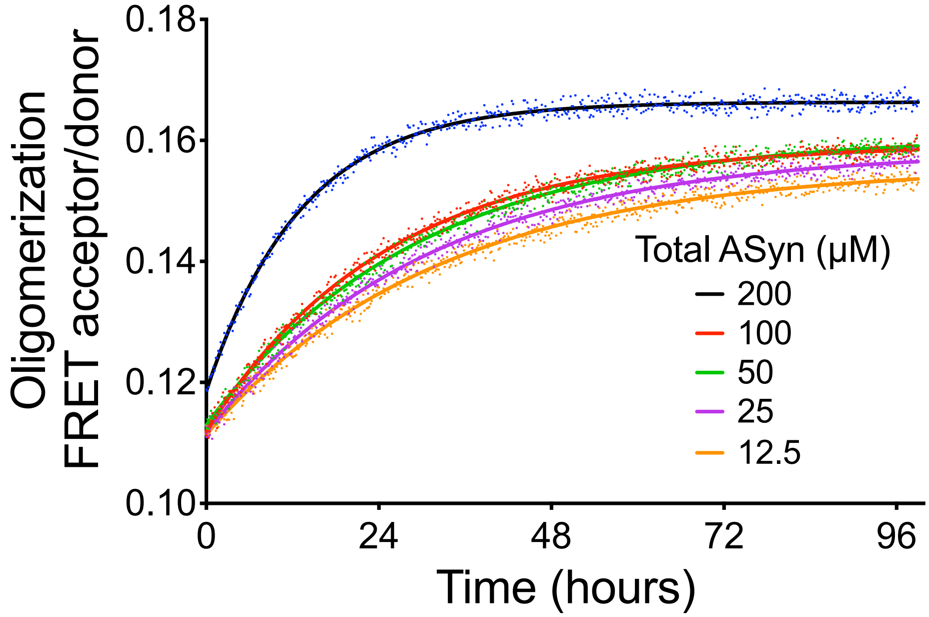


**S4. The dependence of the FRET oligomerization assay on ASyn concentrations**.
A FRET based ASyn oligomerization assay was performed at various concentrations of ASyn as reported[4]. The means of three replicates and first order kinetic fit (line) calculated using Prism software are shown. The initial rates calculated from these data were used to generate Fig 1F in the main manuscript. A subset of these data was shown in a prior publication [4].

### Methods

#### Molecular cloning

PLVSyn, a gift from Brian Spencer, expresses untagged wild-type human ASyn driven from the CMV promoter in the lentivirus vector described[5]. S1 and S2, gifts from Pamela McLean, express ASyn fused to the N and C terminal split fragments of Gaussia luciferase respectively[2]. pLMck1 expression vector was generated by inserting into plasmid pcdna3.1+ the hybrid adenovirus MLP IgG splice sequence shown to enhance expression in mammalian cells[6] at the 5 prime end of the multi cloning site. The following vectors express the indicated proteins in the vector pLck1: pckSyn-SmBit and pckSyn-LgBit express ASyn fused at its C-terminus with a linker followed by the small bit or large bit respectively of split nLuc as described for our bacterial expression vectors. pckMBP-LgBit expresses maltose binding protein fused at its C-terminus with the large bit of the NLuc system[3] with the linker sequence IDGGGGSGGGGSSG between the two proteins. PCK_T1 expresses fluorescent protein mTurquoise2[7]. pCKSynT1 expresses ASyn tagged at its C-terminus with the linker GSAAPVATGS followed by mTurquoise2 coding sequence. Additional plasmids are described in the methods of the main paper.

#### Cellular nLuc assay

H4 cells are plated at $7x10^3$ cells per well in a polyD-Lysine coated clear bottom white 96 well plate in DMEM plus 10% FCS and the next day were transfected with DNA and Fugene (Promega, Madison, WI) as per manufacturer's recommendations using a ratio of transfection reagent to DNA of 4:1. 24 hours after transfection the media was replaced with Opti-MEM without phenol red medium (Opti-MEM). The next day, as a surrogate measure of cell densities, the plates were read on a microplate reader SpectraMax M5 (Molecular Devices LLC,) with an excitation wavelength of 420nm and emission wavelength of 480nm to determine turquoise fluorescence levels. To harvest the media, the plates were spun for 5 minutes at 1050 rpm in an Eppendorf, model 5804 and 40 μl of supernatant was transferred to a 96 well white plate and mixed with 40 μl Opti-MEM. Cells were washed in PBS and lysed in 30 μl of cell lysis buffer (luciferase cell lysis buffer, NEB Catalog # B3321) at room temperature and mixed on an orbital shaker for 30 minutes. Oligomerization was quantified by measuring the luminescence on a SpectraMax L Microplate Reader (Molecular Devices). 80 µl for media or 100 µl for cells of Coelenterazine (CTZ) native substrate (40 µM, NanoLight Technology) in Opti-MEM was injected into the wells with the samples using the injector integrated in SpectraMax L. nLuc activity was measured using 0.1 sec delay after the addition of substrate and 2 sec integration time. Additional cell assay methods are described in the methods of the main paper.

#### Cell staining and Imaging

H4 cells were plated at $7x10^3$ cells per well in DMEM with 10% FCS in a polyD-Lysine coated μclear black 96 well plate (Greiner Bio-One, Monroe, NC) and the next day were transfected with DNA and Fugene (Promega, Madison, WI) as per manufacturer's recommendations using a ratio of transfection reagent to DNA of 4:1. The following day, the media was replaced with fresh media. 72 hours after transfection cells were fixed for 15 minutes in 4% PFA. Cells were stained with 1 µg/ml of purified S129 phosphorylated ASyn specific antibody 11A5[1] overnight at 4°C. They were washed 3 times in PBS and incubated with a goat anti-mouse secondary antibody (Invitrogen) at 1/300 concentration and Hoescht 33342 (Invitrogen) diluted at 1/2000 to stain nuclei for 2 hours at room temperature. Cells were then washed 3 times in PBS. Nuclei and ASyn aggregates were imaged on an Arrayscan XTI high content automated imaging system (Thermo Fisher, Cellomics) and analyzed for aggregates using the spot-detector paradigm of the HCS Studio™ Cell Analysis Software.
